# Implementation of a flavin biosynthesis operon improves extracellular electron transfer in bioengineered Escherichia coli

**DOI:** 10.1101/2022.12.31.522390

**Authors:** Mohammed Mouhib, Melania Reggente, Ardemis A. Boghossian

## Abstract

Bioelectrochemical systems (BES) are promising for energy, sensing, environmental, and synthesis applications. *Escherichia coli* were previously bioengineered for application in BES by introduction of extracellular electron transfer (EET) pathways. Inspired by the metal-reducing (Mtr) pathway of *Shewanella oneidensis* MR-1, several of its cytochromes were heterologously expressed in *E. coli*, leading to increased EET rates and successful application in BES. Besides direct electron transfer*, S. oneidensis* MR-1 is known to secrete flavins that act as redox mediators and are crucial for high EET rates.

Here we co-express the Mtr pathway and a flavin biosynthesis pathway in *E. coli*, to enhance EET in engineered strains. The secretion of both flavin mononucleotide and riboflavin was increased up to 3-fold in engineered strains. Chronoamperometry revealed an up to ~3.4-fold increase in current over the wild type when co-expressing cytochromes and flavin biosynthesis genes, and a ~2.3-fold increase when expressing flavin biosynthesis genes on their own. Thus, the introduction of flavin biosynthesis genes yields in a distinct, yet complementary EET mechanism, and holds promise for application in BES.

## Introduction

Microbes offer a largely untapped potential for the development of sustainable electrochemical systems for energy^[1]^, sensing^[2–4]^, environmental^[3–5]^, and synthesis^[6–8]^ applications. At the core of these technologies lies extracellular electron transfer (EET), which enables the exchange of energy and information between microbes and electrodes in bioelectrochemical systems (BES). The ideal microorganism for a given application should thus be capable of efficient EET, and adaptable to the specific requirements of the application, such as the ability to catabolize certain metabolites, react to environmental stimuli, deactivate pollutants, or produce specific chemicals.

A multitude of microbes was found to be capable of EET^[9]^, and *S. oneidensis* MR-1 has emerged as a notable exoelectrogen due to extensive elucidation of its electron transfer mechanisms and application in BES^[1,10]^. In *S. oneidensis* MR-1, electrons are mainly transferred to extracellular electron acceptors using the Mtr pathway and soluble, as well as cytochrome-associated flavins^[10,11]^. The Mtr pathway consists of an inner membrane quinol oxidase, periplasmic cytochromes, and the outer membrane-spanning Mtr-complex^[10,11]^. Throughout the pathway, electrons are transferred through covalently bound heme cofactors in c-type cytochromes. While the Mtr pathway enables direct electron transfer to an extracellular electron acceptor, *S. oneidensis* MR-1 was shown to also rely on soluble flavins for EET, enabling diffusion-based EET in solution and across thick biofilms^[12–14]^. Indeed, Marsili et. al. could attribute approximately 70% of EET in a *S. oneidensis* MR-1 based electrochemical measurement to flavins secreted to the culture medium over time, with riboflavin being the most prevalent^[12]^.

While these mechanisms equip *Shewanella* species with the efficient EET capabilities necessary for implementation in BES, *S. oneidensis* MR-1 and other native exoelectrogens are often lacking in their substrate spectrum and genetic amenability to realize specific BES applications. Therefore, *E. coli* has emerged as a promising alternative to native exoelectrogens for BES applications, and as a target for engineering to enable efficient EET^[4,6,15–22]^. Over the past two decades, different *S. oneidensis* MR-1 cytochromes were expressed in *E. coli*, ranging from expression of MtrA by Pitts. et. al.^[15]^, overexpression of the full MtrCAB complex in the outer membrane^[16]^ and co-expression of the inner membrane cytochrome CymA^[18]^, to the expression of both membrane-associated and periplasmic cytochromes^[22]^. These advances in implementing direct EET mechanisms in *E. coli* are promising and have led to applications in electrode-assisted biosynthesis of chemicals and biosensing^[4,6,17,20,23]^.

Beyond direct electron transfer through cytochromes, EET in engineered *E. coli* can be further improved using exogenous mediators^[6,17–18,21,23]^. Indeed, commonly available lab strains of *E. coli* are limited in their ability to form biofilms, making it challenging to rely on direct electron transfer in BES with no specific immobilization on the electrode^[24]^. However, exogenous mediators may entail additional costs and technical challenges^[25]^. Endogenous mediators overcome these challenges, and *E. coli* have previously been engineered for secretion of phenazine-1-carboxylic acid by Feng et. al^[26]^.

Since endogenous flavins play a crucial role as redox mediators in EET by *S. oneidensis* MR-1, several studies focused on improving EET targeted to increase their availability. This was achieved by co-culture with flavin-secreting bacteria, or genetic introduction of a flavin biosynthesis module for increased flavin production by *S. oneidensis* MR-1^[27–29]^. Heterologous expression of a flavin biosynthesis pathway from *Bacillus subtilis* in *S. oneidensis* MR-1 by Yang et. al. was shown to impact EET rates in a microbial fuel cell, leading to a 13-fold increase in the maximum power output^[29]^. Biosynthesis of flavins has also been demonstrated using *E. coli* with the goal of developing a riboflavin-producing strain^[30]^. While *E. coli* natively encode genes necessary for flavin biosynthesis, their overexpression or expression of a *Bacillus subtilis* pathway can greatly increase secreted flavin concentrations^[30]^.

Based on previous findings on the important role of flavins in *S. oneidensis* MR-1^[12]^, we aimed to explore the biosynthesis of riboflavin and flavin mononucleotide to complement the Mtr pathway in exoelectrogenic *E. coli* (Figure 1a). Therefore, we equipped *E. coli* C43 (DE3) with both the Mtr pathway and a synthetic operon for flavin biosynthesis. Obtained strains were characterized for flavin secretion profiles, as well as cytochrome expression and localization. Finally, we evaluated EET using chronoamperometry in a three-electrode single chambered electrochemical setup.

**Figure 1.**
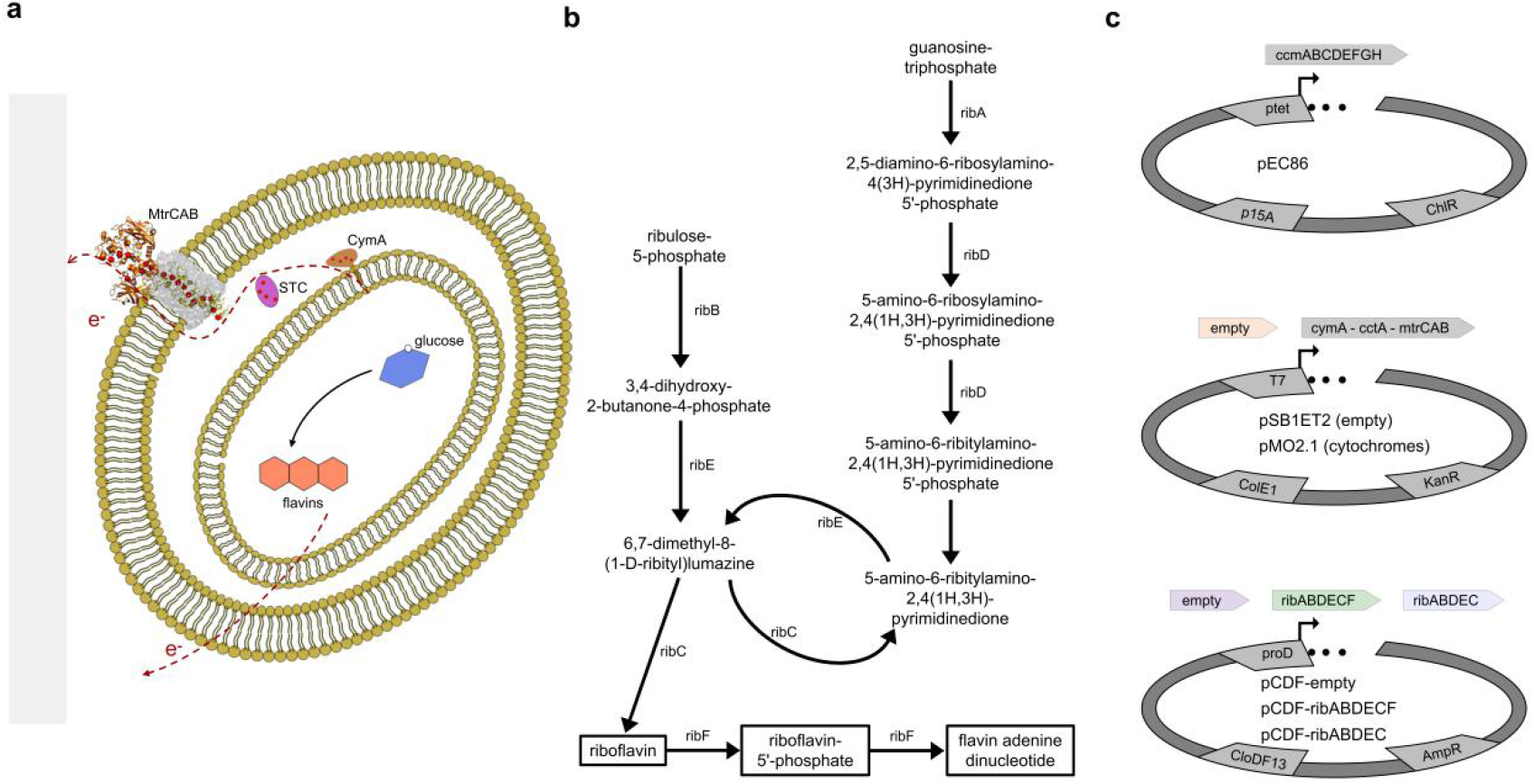
Co-expression of the Mtr pathway and a flavin biosynthesis pathway. (a) Schematic of an *E. coli* cell expressing both a flavin biosynthesis pathway and the Mtr pathway across the inner membrane, periplasm and outer membrane. (Not to scale) (b) Overview over the flavin biosynthesis pathway expressed during this study. Genes encoding enzymes catalyzing a given reaction are denominated next to arrows connecting metabolites. (c) Illustration of plasmids used in this study, including the promoters (ptet, T7, proD), origins of replication (p15A, ColE1, CloDF13), and antibiotic resistance genes (ChlR - chloramphenicol, KanR - kanamycin, AmpR - ampicillin). Expressed genes of interest are illustrated by arrows above the corresponding plasmids.

## Results and discussion

For the secretion of flavins as redox mediators, we leveraged *E. coli*’s native ability for biosynthesis of riboflavin and flavin mononucleotide (Figure 1b)^[30]^. Collectively, the GTP cyclohydrolase II (*ribA*), 3,4-dihydroxy-2-butanone 4-phosphate synthase (*ribB*), fused diaminohydroxyphosphoribosylaminopyrimidine deaminase / 5-amino-6-(5-phosphoribosylamino)uracil reductase (*ribD*), 6,7-dimethyl-8-ribityllumazine synthase (*ribE*) and riboflavin synthase (*ribC*) enable biosynthesis of riboflavin from guanosine triphosphate and ribulose-5 phosphate. Further reactions of riboflavin to riboflavin-5-phosphate (FMN) and flavin adenine dinucleotide are catalysed by a bifunctional riboflavin kinase / FMN adenylyltransferase^[30,31]^. The encoding genes, *ribABDECF*, are not organized in one operon on the chromosome^[31]^. Therefore, we opted to construct a synthetic operon for the expression of either *ribABDEC* to accumulate riboflavin, or *ribABDECF* to accumulate FMN, under the control of a strong constitutive promoter^[32]^. In addition to a plasmid enabling flavin biosynthesis, we also employed a plasmid for expression of the Mtr pathway^[22]^ and a plasmid encoding the ccm machinery^[33]^ necessary for post-translational modification of c-type cytochromes, yielding a total of five strains (Figure 1c, Table 1).

**Table 1.** Strain designations by transformed plasmid DNA

| 1 | 2 | 3 | name |
| --- | --- | --- | --- |
| pEC86 | pSB1ET2 | pCDF-empty | empty |
|  |  | pCDF-*ribABDEC* | *rib-dF-dCyt* |
|  | pMO2.1 | pCDF-empty | *cymstcmtr* |
|  |  | pCDF-*ribABDECF* | *rib* |
|  |  | pCDF-*ribABDEC* | *rib-dF* |

To evaluate whether the Mtr pathway is expressed and flavins are secreted when co-expressing both constructs, we performed aerobic growth experiments in a minimal medium with glucose as the sole carbon source. After cytochrome expression in strains with either no flavin biosynthesis module (cymstcmtr), co-expression of *ribABDECF* genes (rib) or co-expression of the *ribABDEC* genes (rib-dF), we followed bacterial growth (Figure S1), as well as the concentrations of riboflavin and FMN over time (Figures 2a and b). While all strains reached comparable optical densities and released both riboflavin and FMN over the course of 48 hours, strains expressing either one of the flavin modules secreted significantly higher amounts of FMN (Figures 2a and b). In addition, the rib-dF strain also showed a significant increase in riboflavin concentration throughout the measurement. After 48 hours the rib strain accumulated the highest total flavin concentration (riboflavin + FMN, Figure 2c) at 10.4 ± 0.61 μM, representing a 20% increase over the rib-dF strain (8.7 ± 0.85 μM), and a three-fold increase over the cymstcmtr strain (3.07 ±0.1 μM). Most of the increase can be attributed to FMN, with only a 24% increase in the riboflavin concentration secreted from the rib strain over the cymstcmtr strain. In contrast, the higher flavin secretion from the rib-dF strain over the cymstcmtr strain represents an approximately 2.4- and 3.3-fold increase in the riboflavin and FMN concentrations respectively. This discrepancy in the FMN to riboflavin (RF) ratio (Figure 2c) was expected, as deletion of the *ribF* gene from the *ribABDECF* operon reduces the amount of available riboflavin kinase for phosphorylation of riboflavin to FMN.

**Figure 2.**
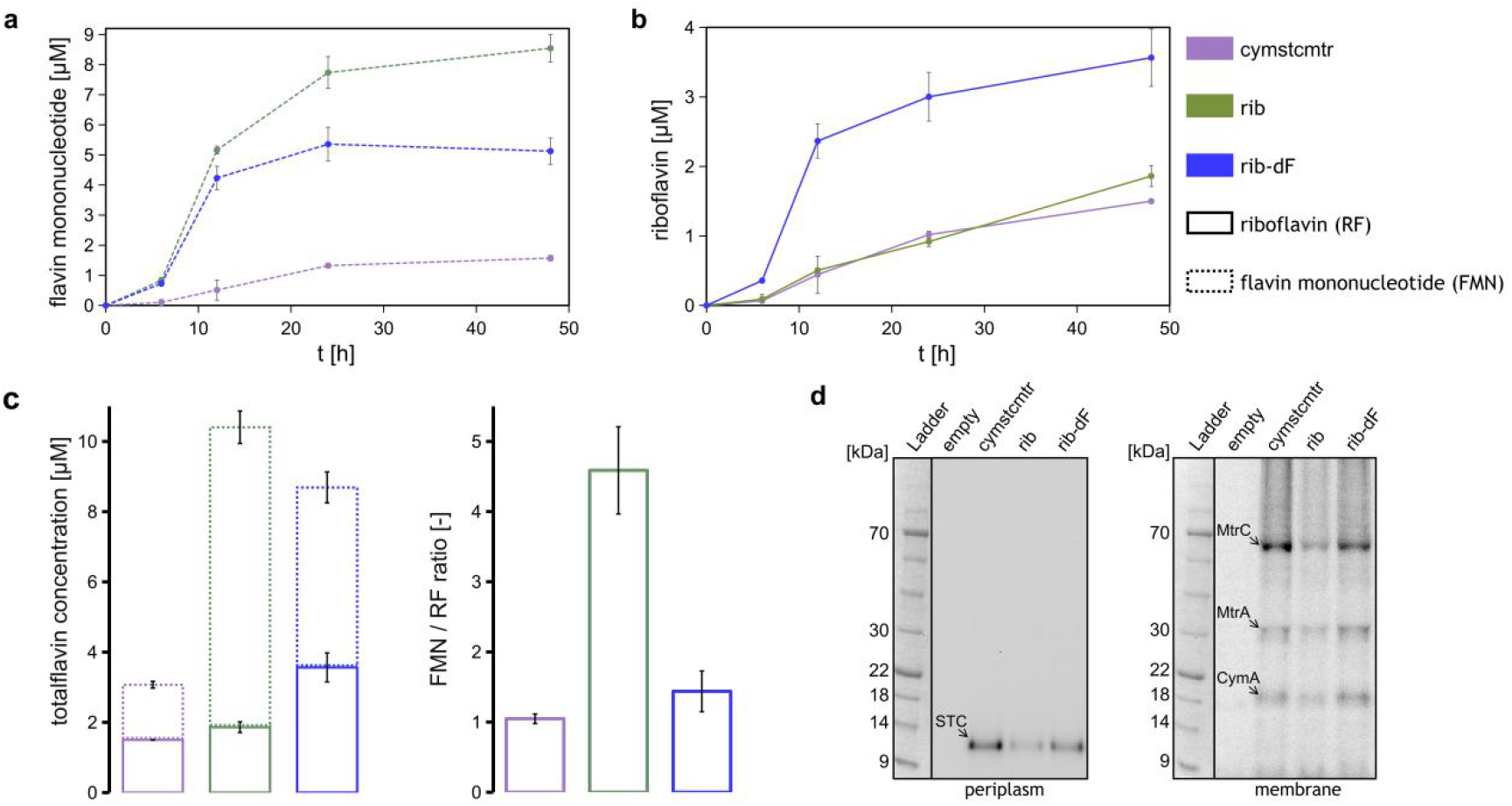
Flavin secretion and cytochrome expression. The concentrations of (a) FMN and (b) riboflavin increase over time during aerobic growth in M9 glucose medium. (c) Total flavin concentration made up of riboflavin and FMN (left), and the ratio of FMN to riboflavin (right) after 48 h of aerobic growth. (c) SDS-PAGE gels (4%-20%, MOPS-SDS buffer, 30 μg of protein per lane) loaded with periplasmic (left) and membrane (right) protein extracts were stained for hemes using an enhanced chemiluminescence substrate. Mean values of 3 independent biological replicates are plotted, with error bars representing 1 standard deviation.

In order to assess correct localization and post-translational modification of Mtr pathway cytochromes under concomitant overexpression of flavin biosynthesis genes, we collected subcellular fractions of modified *E. coli* and used ECL staining to identify full-length c-type cytochromes in resulting protein samples. All strains expressed the small tetraheme cytochrome STC (*cctA* gene product) in the periplasm, as well as the inner membrane cytochrome CymA and components of the Mtr complex (MtrA, MtrC) in the membrane fraction (Figure 2d). The protein concentrations in periplasmic and membrane extracts were kept constant between samples from different strains, and controlled by total protein staining (Figure S2). At equal loading, cytochrome concentrations appear to be higher when no riboflavin biosynthesis genes are over-expressed (cymmtr), as compared to cytochrome expression in the rib and rib-dF strains (Figure 2d). This trend was validated using further replicates (Figure S3), revealing that in addition to higher riboflavin concentrations, the rib-dF strain also showed an increased cytochrome expression as compared to the rib strain (Figures 2d, S3).

To evaluate whether flavin biosynthesis may enhance EET in engineered *E. coli*, we performed chronoamperometry measurements under anaerobic conditions with glucose as the sole carbon source, and an electrode kept at +0.2 V against an Ag/AgCl reference as the electron acceptor. While both the rib, and rib-dF strains showed increases in current over the empty vector control, the rib-dF strain achieved significantly higher currents than the rib strain (Figure S4). Therefore, further electrochemical characterization was focused on assessing whether increased EET in the rib-dF strain is comparable with the sole expression of either the Mtr pathway or riboflavin biosynthesis genes (Figure 3a). Surprisingly, the sole expression of the *ribABDEC* genes (rib-dF-dCyt) led to currents on par with the expression of the Mtr-pathway. Biosynthesis and secretion of flavins thus offer a distinct way to improve EET in engineered *E. coli*. Co-expression of both pathways led to ~3.4-fold current increase over the wild type, and a ~50% increase over the cymstcmtr and *ribABDEC* strain respectively. While flavin-secreting strains rib-dF, rib, and rib-dF-dCyt showed increased EET, we observed declining optical densities over the course of chronoamperometry (Figures 3b, S4). In contrast, both the empty and cymstcmtr strains showed growth over the same time frame, suggesting that overexpression of flavin biosynthesis genes negatively impacts strain viability.

**Figure 3.**
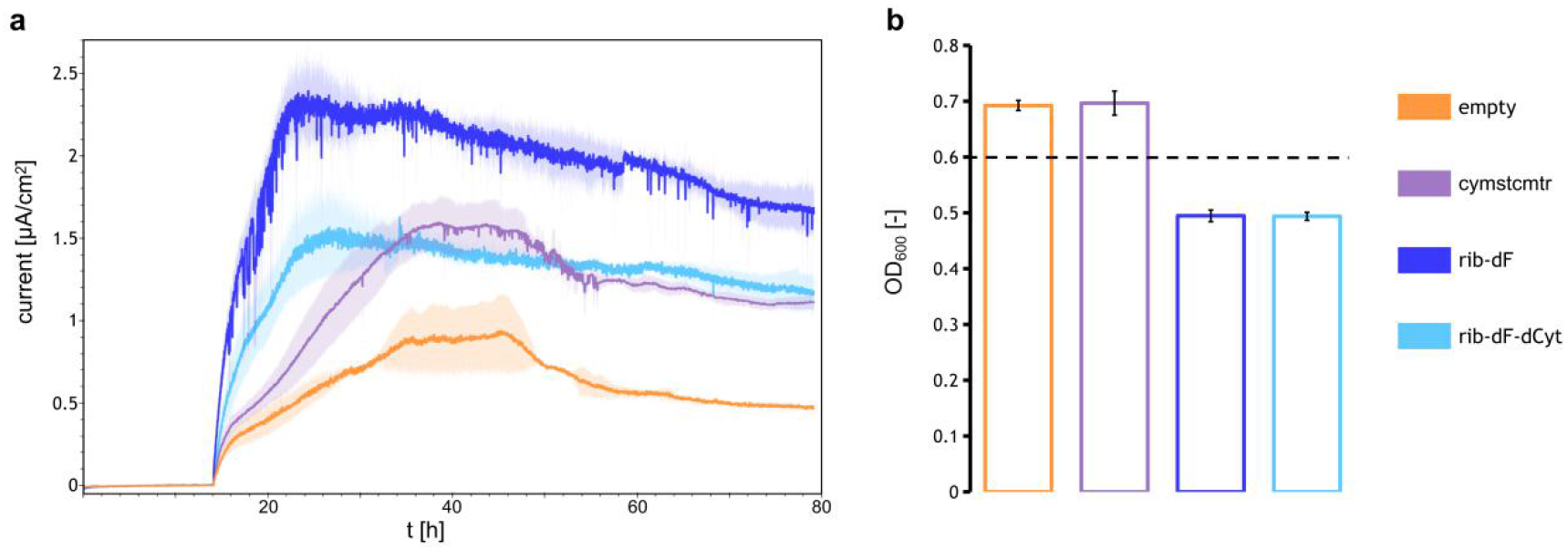
EET characterization using grahite felt as an electron acceptor. (a) Chronoamperometric measurements under anaerobic conditions in M9-glucose medium. A positive potential of 0.2 V against an Ag/AgCl reference electrode was applied, and current values were recorded every 30 seconds. Mean currents of three independent measurements are plotted over time (two measurements for the empty vector control), with the shaded area representing one standard deviation. (b) The OD_600_ in solution was measured after chronoamperometry, starting at an OD_600_ of 0.6 (black line). Bars represent mean values, with the error given as one standard deviation.

## Conclusion

We assessed flavin biosynthesis in the cymstcmtr, rib and rib-dF strains under aerobic conditions as well as cytochrome expression and localization. Both riboflavin and FMN were produced in higher concentration after introduction of the respective biosynthesis pathways, with the highest riboflavin concentration achieved in the rib-dF strain. All cytochromes of the Mtr pathway were expressed and correctly localized across strains, in agreement with previous expression in the absence of flavin biosynthesis gene overexpression^[22]^. However, expression levels decreased for both flavin producers, with the rib strain showing the lowest cytochrome expression.

Based on the importance of flavins as redox mediators in *S. oneidensis* MR-1^[12]^ and previous engineering efforts to further increase its flavin secretion^[29]^, EET was expected to be enhanced in the presence of flavin biosynthesis genes. Indeed, we observed substantially increased currents during chronoamperometry, with the rib-dF strain showing the highest EET rate with a ~3.4-fold current increase over the control strain (empty). It is notable that an increase in current was also observed for sole expression of the riboflavin biosynthesis genes *ribABDEC*, showing that flavin biosynthesis offers a distinct EET pathway which is not dependent on Mtr pathway cytochromes. While sole expression of cytochromes and a flavin biosynthesis pathway enhance EET separately, co-expression still yielded the highest EET rates. Thus, future work should focus on carefully balancing expression levels between these two components. This may improve strain fitness and could lead to retained cytochrome expression levels under concomitant overexpression of a flavin biosynthesis pathway.

Both cytochrome expression and flavin biosynthesis offer independently functional, and complementary ways to improve EET in *E. coli*. This modularity holds promise in furthering EET rate enhancements in *E. coli* through fine tuning of expression levels, as well as in implementation in energy, sensing, environmental and synthesis applications.

## Experimental section

### Molecular cloning

Plasmids, primers, and strains used in this study can be found in the supplementary information (Table S1). The empty vector pCDF-empty was cloned using Gibson assembly^[35]^. The backbone was amplified from pCDF-mcherry1 and the ampicillin resistance gene was amplified from a pET vector using primers gibson_amppCDF_fwd, gibson_amppCDF_rev, gibson_ampR_fwd and gibson_ampR_rev. The genes *ribA*, *ribB*, *ribDE* and *ribF* were all amplified from an *E. coli* BL21 (DE3) genome extract using the corresponding primers in Table S1. A total of six fragments was assembled using splicing by overlap extension PCR (primers SOE1_fwd, SOE1_rev, SOE2_fwd, SOE2_rev) in two separate reactions (SOE1: *ribDE*, *ribC*, *ribF;* SOE2: pCDF, *ribA*, *ribB*). The final products (SOE1 and SOE2) were joined using Gibson assembly to yield plasmid pCDF-ribABDECF. For cloning of the plasmid pCDF-ribABDEC, pCDF-ribABDECF was amplified using primers d_ribF_fwd and d_ribF_rev, followed by blunt end ligation using T4 DNA ligase. All cloning steps were performed using *E. coli* Dh5α, and all PCR reactions were performed using Q5 DNA polymerase (NEB).

### Cytochrome expression

For cytochrome expression, three plasmids were co-transformed into electrocompetent *E. coli* C43 (DE3) using different combinations outlined in Table 1, to obtain the strains, emtpy”, "rib-dF-dCyt”, cymstcmtr”, "rib”, and "rib-dF”. For selection, kanamycin (50 μg/ml), chloramphenicol (34 μg/ml), and carbenicillin (100 μg/ml) were used in both agar plates and liquid 2x YT medium. Over night cultures were grown from single colonies in 5 ml 2x YT medium (15 ml tubes) at 37°C and 220 rpm shaking. Expression cultures (25 ml 2xYT medium in 250 ml baffled Erlenmeyer flasks, 220 rpm shaking, supplemented with 1 mM 5-aminolevulinic acid) were inoculated to an OD_600_ of 0.1 from over night cultures, grown to an OD_600_ of approximately 0.6 at 37°C, followed by induction using 10 μM IPTG and over night growth at 30°C for 16 h.

### Subcellular fractionation

Subcellular fractionation was carried out exactly as previously described^[22]^. For recovery of periplamic fractions, bacteria were centrifuged, washed in PBS, resuspended in buffer A (100 mM Tris-HCl pH 8, 500 mM sucrose, 0.5 mM EDTA), incubated for 5 minutes on ice, recentrifuged and resuspended (gently using an inoculation loop) in 1 mM MgCl_2_ followed by a 2 min incubation step on ice. After a final centrifugation step, supernatants were collected as the periplasmic fraction, and concentrated using 3 kDa molecular weight cut-off Amicon filters. All centrifugation steps for recovery of periplasmic protein were carried out at 4°C and 3000 g for 15 minutes per step.

Pellets were used for recovery of membrane proteins. Bacteria were washed in buffer B (50 mM Tris-HCl pH 8, 250 mM sucrose, 10 mM MgSO4), resuspended in buffer C (50 mM Tris-HCl pH 8, 2.5 mM EDTA) and lysed using sonication. Lysates were cleared by centrifugation (10000 g, 4°C, 10 min), and membrane proteins were pelleted from lysates by centrifugation at 21000 g for 4 h at 4°C. Membrane pellets were washed once in buffer C, re-centrifuged, and finally resuspended in buffer C supplemented with 1% Triton X-100.

### SDS-PAGE and cytochrome staining

Protein samples were separated at 150 V in 4% - 20% SurePage (Genscript) polyacrylamide gels using MOPS-SDS running buffer and 1x NuPAGE LDS sample buffer. After SDS-PAGE, gels were rinsed in water followed by light drying to remove excess water using a tissue. After application of an ECL stain (SuperSignal West Pico PLUS, Thermo Scientific) and incubation for 5 minutes, gels were imaged using an ECL imager (Fusion solo S, Vilber)

### Electrochemical setup and chronoamperometry

Chronoamperometry was carried out in a single chambered 3 electrode setup, with an applied potential of 0.2 V against an Ag/AgCl reference electrode. The anode was composed of a platinum wire connected to a 1 x 1.1 x 0.5 cm piece of graphite felt (Alfa Aesar), and a platinum wire served as the cathode. The felt was pre-treated by immersion in a piranha solution (97% H2SO4 : 30% H2O2 in 3:1 mixture) for 10 minutes, followed by washing and storage in distilled water. Chambers were filled with 95 ml M9-medium, prepared according to Baruch et. al.^[40]^ and supplemented with kanamycin (50 μg/ml), IPTG (10 μM) and glucose (0.4%). Cytochrome expression cultures were washed (3x 10 ml; 4°C, 3000g, 20 min centrifugation) and resuspended (5 ml) in the same medium, followed by injection into the electrochemical setup during chronoamperometry (final volume of 100 ml) to a final OD_600_ of 0.6. Following chronoamperometry, the OD_600_ was measured to assess bacterial growth. Anaerobic conditions were set using a constant stream of N_2_(g) throughout the experiment.

### HPLC determination of flavins

After cytochrome expression in 2x YT medium (see cytochrome expression), cultures were centrifuged (4°C, 3000 g, 20 min) and pellets washed in M9-glucose medium (see electrochemical setup and chronoamperometry). After 3 washes in 10 ml M9-glucose medium each, bacteria were used to inoculate 10 ml of M9 glucose medium in 100 ml Erlenmeyer flasks, supplemented with kanamycin (50 μg/ml), chloramphenicol (34 μg/ml) and carbenicillin (100 μg/ml). Cultures were kept at 30°C and 220 rpm shaking with 800 μl samples taken after 0 h, 6 h, 12 h, 24 h, and 48 h.

Growth was monitored using OD_600_ measurements on a spectrophotometer (UV-3600 Plus, Shimadzu). Filtered samples (0.2 μm syringe filters) were used for HPLC measurements (Agilent Infinity II). Samples were separated on a C18 column (Agilent ZORBAX Eclipse Plus C18, RRHD, 2.1 x 50 mm, 1.8 μm) using ammonium acetate buffer (pH 6) and methanol as the mobile phase (0.9 ml/min flow rate, 30°C). The methanol concentration in the mobile phase was kept at 8% for 1 minute, increased to 20% over 7 minutes, increased to 70% over 30 seconds, kept at 70% for 3 minutes, decreased to 8% over 30 seconds, and finally kept at 8% for 3 minutes (15 minutes total run-time per sample). Absorbance was measured at 210 nm, 270 nm, 375 nm and 450 nm (Agilent 1260 DAD WR).

## Conflicts of interest

There are no conflicts to declare.

## Acknowledgements

The authors are grateful for funding by the Gebert-Rüf Stiftung.

## Notes

### Competing Interest Statement

The authors have declared no competing interest.

