## supplementary_information for "Implementation of a flavin biosynthesis operon improves extracellular electron transfer in bioengineered Escherichia coli"

**Table S1. Overview of bacterial strains, plasmids and primers used in this study.**

| Strains, plasmids and primers | Characteristics | Source |
|---|---|---|
| **Strains** | | |
| *E. coli* DH5α | | |
| *E. coli* C43 (DE3) | | |
| **Plasmids** | | |
| pEC86 | CcmABCDEFGH expression, chloramphenicol resistance, tet promoter | [33] |
| pSB1ET2 | empty backbone for cytochrome expression, kanamycin resistance, T7 promoter | [16] |
| pMO2.1 | CymA, STC and MtrCAB expression, kanamycin resistance, T7 promoter | [22] |
| pCDF-mcherry | constitutive mcherry expression, spectinomycin resistance, proD promoter | *** |
| pCDF-empty | empty vector for flavin biosynthesis genes, ampicillin resistance, proD promoter | this study |
| pCDF-ribABDECF | ribABDECF encoding plasmid, ampicillin resistance, proD promoter | this study |
| pCDF-ribABDEC | ribABDEC encoding plasmid, ampicillin resistance, proD promoter | this study |
| **Primers (5`- 3`)** | | |
| gibson_ribpCDF_fwd | CCGCCCGCGAATTTTTTGGGCTAACAAAACCGGCTTAATAAGGACGAGCCTCAGACC | |
| gibson_ribpCDF_rev | AGTTTGGCTTCTGCCACACGTTTAAGCTGCATCTAGTATTTCTCCTCTTTCTCTAGTAGC | |
| gibson_ribA_fwd | GATCGCATGGTTGCTACTAGAGAAAGAGGAGAAATACTAGATGCAGCTTAAACGTGTGG | |
| gibson_ribA_rev | GTAAAAAAACCTCACTGAAATTATGGTTACCAGAATCAGCAAGAGGGTTATTTGTTCAGC | |
| gibson_ribB_fwd | TGGGCCATTTGCTGAACAAATAACCCTCTTGCTGATTCTGGTAACCATAATTTCAGTGAG | |
| gibson_ribB_rev | CTCCAGGCGCGCGATCTCTTCGCCAAATTCTTTAAGCAGCGGTTTTCAGCTG | |
| gibson_ribDE_fwd | CACATGAGCGTAAAGCCAGCTGAAAACCGCTGCTTAAAGAATTTGGCGAAGAGATCG | |
| gibson_ribDE_rev | CAACTCCTGAAATCAGTTAAGACATTCTGTTCAGTTACTAATTTCAGGCCTTGATGG | |
| gibson_ribC_fwd | TTGAAAGCCATCAAGGCCTGAAATTAGTAACTGAACAGAATGTCTTAACTGATTTCAGG | |
| gibson_ribC_rev | TCTGGCTCAAAACAGTGAAAATCGTCCGAGTAGATTTCAGATCAGGCTTCTGTACC | |
| gibson_ribF_fwd | ATCAACCAGGTACAGAAGCCTGATCTGAAATCTACTCGGACGATTTTCACTGTTTTGAG | |
| gibson_ribF_rev | GTCAGGTATGATTTAAATGGTCTGAGGCTCGTCCTTATTAAGCCGGTTTTGTTAGCCCA | |
| SOE1_fwd | CCGCCCGCGAATTTTTTGGGCTAAC | |
| SOE1_rev | CTCCAGGCGCGCGATCTCTTCG | |
| SOE2_fwd | CACATGAGCGTAAAGCCAGCTG | |
| SOE2_rev | GTCAGGTATGATTTAAATGGTCTGAGGCTC | |
| gibson_amppCDF_fwd | ACTCTTCCTTTTTCAATATTATTGAAGCATTTATCAGG | |
| gibson_amppCDF_rev | GATAGGTGCCTCACTGATTAAGCATTGGTAATGTCTAACAATTCGTTCAAGCCGAG | |
| gibson_ampR_fwd | TCGGCTTGAACGAATTGTTAGACATTACCAATGCTTAATCAGTGAGGCAC | |
| gibson_ampR_rev | GATAAATGCTTCAATAATATTGAAAAAGGAAGAGTATGAG | |
| d_ribF_fwd | TAA GGA CGA GCC TCA GAC C | |
| d_ribF_rev | AGA TCA GGC TTC TGT ACC TGG | |

***pCDF-mcherry1 was a gift from Michael Lynch (Addgene plasmid # 87144 ; http://n2t.net/addgene:87144 ; RRID:Addgene_87144)

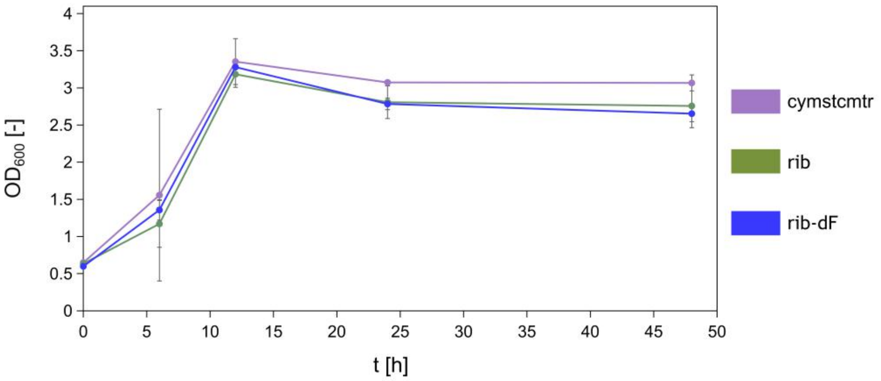


**Figure S1.** Change in $OD_{600}$ over time during aerobic growth in M9-glucose medium, starting at an $OD_{600}$ of 0.6.

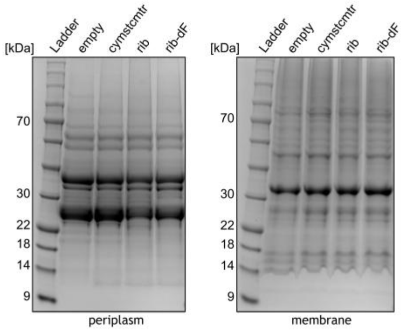


**Figure S2. Total protein staining of SDS-PAGE gels.** Gels were stained using Coomassie brilliant blue G250 following prior ECL staining of the same gels (see Figure 2d).

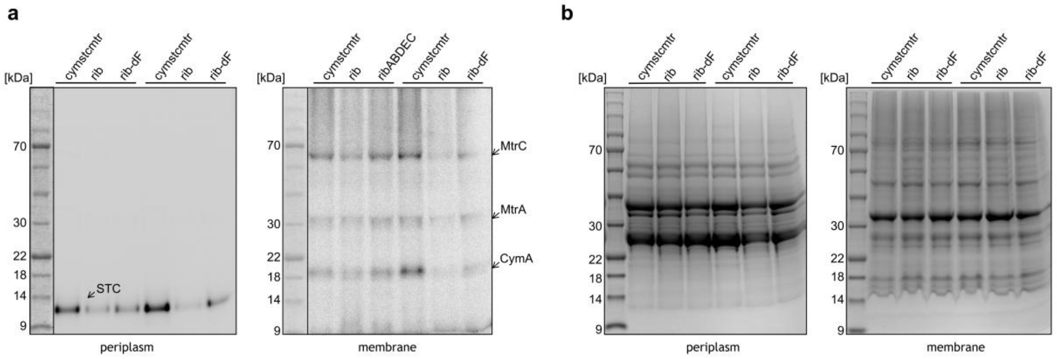


**Figure S3. Addittional replicates to assess cytochrome expression and localiztion.** SDS-PAGE gels (4%-20%, MOPS-SDS buffer, 30 µg of protein per lane) loaded with periplasmic and membrane protein extracts were stained for (a) hemes using an enhanced chemiluminescence substrate and (b) total protein content using Coomassie brilliant blue G250.

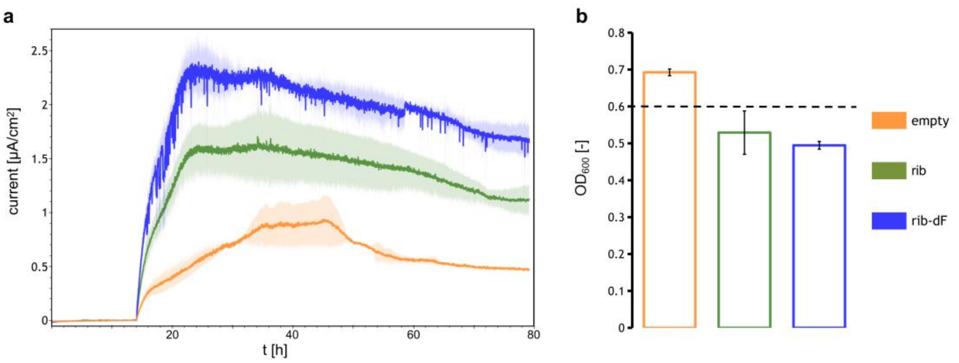


**Figure S4.** (a) Chronoamperometric measurements under anaerobic conditions in M9-glucose medium. A positive potential of 0.2 V against an Ag/AgCl reference electrode was applied, and current values were averaged every 30 seconds. Mean currents of three independant measurements are plotted over time (two measurements for the empty vector control), with the shaded area representing one standard deviation. (b) The $OD_{600}$ in solution was measured after chronoamperometry, starting at an $OD_{600}$ of 0.6 (black line). Bars represent mean values, with the error given as one standard deviation.
